# Advancing the Indian Cattle Pangenome: Characterizing Non-Reference Sequences in *Bos indicus*

**DOI:** 10.1101/2024.07.18.604025

**Authors:** Sarwar Azam, Abhisek Sahu, Naveen Kumar Pandey, Mahesh Neupane, Curtis P Van Tassel, Benjamin D Rosen, Ravi Kumar Gandham, Subha Narayan Rath, Subeer S Majumdar

**Affiliations:** National Institute of Animal Biotechnology, Hyderabad, India; Indian Institute of Technology Hyderabad, Sangareddy, India; Animal Genomics and Improvement Laboratory, USDA-ARS, Beltsville, MD 20705, USA

**Keywords:** Pangenome, *Bos indicus*, Cattle, Linked-reads, Genome assembly, NUIs, BICIs

## Abstract

**Background:** India, with the world’s largest cattle population and more than 50 registered breeds of *Bos indicus*, stands as a vital reservoir of genetic diversity. However, the abundant diversity among Indian cattle breeds highlights the inadequacy of a single reference sequence to represent the entire genomic content of desi cattle. We recognize the need to capture the genomic differences within the *Bos indicus* population as a whole, and specifically within the dairy cattle subset by identifying non-reference sequences and constructing a pangenome.

**Finding:** Five representative genomes of prominent dairy breeds, including Gir, Kankrej, Tharparkar, Sahiwal, and Red Sindhi, were sequenced using 10X Genomics ’Linked-Read’ technology. Assemblies generated from these linked-reads ranged from 2.70 Gb to 2.77 Gb,comparable to the *Bos indicus* Brahman reference genome. A pangenome of *Bos indicus* cattle was constructed by comparing the newly assembled genomes with the reference using alignment and graph-based methods, revealing 8 Mb and 17.7 Mb of novel sequence respectively. A confident set of 6,844 Non-Reference Unique Insertions (NUIs) spanning 7.57 Mbs was identified through both methods, representing the pangenome of Indian *Bos indicus* breeds. Comparative analysis with previously published pangenomes unveiled 2.8 Mb (37%) commonality with the Chinese indicine pangenome and only 1% commonality with the *Bos taurus* pangenome. Among these, 2,312 NUIs - encompassing ∼2 Mb, were commonly found in 98 samples of the 5 breeds and designated as *Bos indicus* Common Insertions (BICIs) in the population. Furthermore, 926 BICIs were identified within 682 protein-coding genes, 54 long non-coding RNAs (LncRNA), and 18 pseudogenes. These protein-coding genes were enriched for functions such as chemical synaptic transmission, cell junction organization, cell-cell adhesion, and cell morphogenesis. The protein-coding genes were found in various prominent Quantitative Trait Loci (QTL) regions, suggesting potential roles of BICIs in traits related to milk production, reproduction, exterior, health, meat, and carcass. Notably, 63.21% of the bases within the BICIs call set contained interspersed repeats, predominantly LINEs. Additionally,70.28% of BICIs are shared with other domesticated and wild species, highlighting their evolutionary significance.

**Conclusion:** This is the first report unveiling a robust set of NUIs defining the pangenome of *Bos indicus* breeds of India. The analyses contribute valuable insights into the genomic landscape of desi cattle breeds.

## Introduction

Cattle hold incredible importance in global agriculture as one of the most pivotal livestock species. They contribute significantly to human nutrition, the economy, and agricultural practices by providing essential resources like milk, meat, hide, and drought power [1,2]. In the Indian context, cattle are predominantly reared for milk production and draught purposes. Cattle can be broadly categorized into two primary types: *Bos taurus* and *Bos indicus*, originating from distinct domestication events [3]. Approximately 10,000 years ago, the Fertile Crescent witnessed the emergence of humpless taurine cattle (*Bos taurus taurus*) [4], while about 8,000 years ago, the Indus Valley gave rise to humped indicine cattle (*Bos taurus indicus*)[5,6]. Genetic studies indicate that these two lineages diverged from a common ancestor approximately 210,000–350,000 years ago, well before the domestication processes took place [5]. This deep genetic divergence underscores the inherent distinctiveness of the ancestral aurochs populations from which both lineages originated. Moreover, multiple migration waves [7], interbreeding, and introgressions with other bovids, such as yak and banteng, have substantially augmented the genetic diversity within the Bovine group [3,8]. Furthermore, the continuous selection and adaptation to diverse climates and environmental pressures, including factors such as altitude and endemic diseases, have further molded and diversified the cattle genome [9]. This has resulted in exceptionally high levels of genetic diversity in cattle populations worldwide.

These genetic differences manifest as unique sequences within the genomes of various cattle breeds. This genomic variation is not limited to single-nucleotide polymorphisms (SNPs) and insertions/deletions (indels) but extends to encompass other structural variations (SVs) [10]. To explore this diversity, initiatives like the 1000 Bull Genomes Project have examined numerous *Bos taurus* breeds, revealing variations that are not universally present in all individuals of the species [11]. Comparisons among various genome assemblies have revealed specific sequences not universally present in all individuals of the species. While a single reference genome was initially considered sufficient to represent the entire species, it later became evident that there are sequences specific to individual breeds within a species [12]. This realization led to the concept of the “pangenome” initially applied to bacterial genomes [13] and later extended to fungi [14] and plant genomes [15]. Surprisingly, this approach had not been extensively explored in large eukaryotes, especially mammals, until the African Pan-genome Project revealed sequence variability in African human genomes compared to the human reference genome [16]. Subsequent reports on the human pangenome established the existence of specific sequences in populations, augmenting the reference genome to construct a comprehensive pangenome [17]. Similar efforts to explore pangenomes were initiated in other large animals, including cattle [12,18,19].

Initially, a read-depth based approach was applied to identify SVs, followed by pangenome construction from these SVs. However, with the availability of multiple genome assemblies, direct sequence comparisons have become more common than mapping reads to the reference genome to identify unique insertions [20]. The latter approach offers a more faithful representation of complex regions and SVs, given the intricacies of calling and representing nested variations. A graph-based method has evolved to compare genome assemblies for the identification of unique insertions, which are then integrated with the reference genome [21]. However, in the context of cattle, only a few efforts have been made to explore the pangenome, primarily within the *Bos taurus* species [12,18,19].

For instance, Zhou et al. [12] reported SVs in the ARS-UCD1.2 *Bos taurus* reference genome [22], using 898 individuals, identifying 83 Mb of sequence not found in the reference genome. Leonard et al. [18] used a trio-binning approach to assemble six genomes, including three *Bos taurus taurus*, one *Bos gaurus*, one *Bos taurus indicus* from hybrid progenies, resulting in the discovery of 90 thousand structural variants, including 931 overlapping with coding sequences. Crysnanto et al. [23] revealed a bovine pangenome using the Hereford-based *Bos taurus* reference genome and five reference-quality assemblies from three taurine cattle breeds (Angus, Highland, and Original Braunvieh) and their close relatives Brahman (*Bos taurus indicus*) and yak (*Bos grunniens*). The pangenome contained an additional 70,329,827 bases compared to the *Bos taurus* reference genome. Their multi-assembly approach unveiled 30 and 10.1 million bases unique to yak and indicine cattle, respectively, as well as between 3.3 and 4.4 million bases unique to each taurine assembly.

While the concept of a pangenome is not limited to specific groups of individuals, populations, or species, there are efforts to create larger pangenomes that encompass multiple species. These expanded pangenomes, sometimes referred to as ’super pangenomes’ [24] have been developed for crops like soybeans [25] and tomatoes [26] , which include both wild and domesticated species. Similar endeavors are being undertaken for cattle by the Bovine Pangenome Consortium [27], which aims to include many wild relatives of Bos. However, it’s important to note that in human and other species, most experiments focus on creating species-specific pangenomes. These species-specific pangenomes are essential and have been instrumental in studying population-specific traits, including disease susceptibility and adaptation.

Linked-Read technology [28] has been successfully employed for developing pangenomes in humans [29]. Wong et al. [29] published a Non-Reference Unique Insertions (NUI) discovery pipeline, identifying missing sequences in the human reference genome from in-silico, phased, de novo human genome assemblies generated using Linked-Reads. They coined the term ’NUI’ to describe unique sequences not present in the reference genome but found in other individual’s genomes. In this study, we present an effort to construct a pangenome specific to Indian *Bos indicus* cattle. In fact, there are more than 50 registered indigenous breeds of *Bos indicus* cattle in India, of which only a few are dairy breeds. The prominent dairy breeds include Gir, Kankrej, Tharparkar, Sahiwal, and Red Sindhi [30,31] . Representative genomes of these five *Bos indicus* breeds were sequenced using Linked-Reads. The assemblies generated were compared with the Brahman genome assembly as currently recognized as the *Bos indicus* reference sequence [32]. Apart from applying the NUI pipeline, we used a graph-based pangenome method to identify unique insertions across *Bos indicus* breeds. Importantly, our effort has led to the development of the first pangenome specific for *Bos indicus* dairy breeds found in India.

## Results

### Sequencing of desi cattle using 10X linked-reads and genome assembly

Genome sequencing of five indigenous cattle breeds, namely Gir, Tharparkar, Kankrej, Sahiwal, and Red Sindhi, was carried out using the 10X Chromium technology [33]. Each genome was sequenced at approximately 100x coverage with 150bp paired-end reads. Assemblies ranged in size from 2.70 Gb to 2.77 Gb for different breeds (**Table 1**). Notably, the Sahiwal and Red Sindhi assemblies displayed fewer contigs and larger N50 values compared to Gir, Kankrej, and Tharparkar. In particular, the Sahiwal assembly featured the largest contig, measuring 156 Mb, while the largest contigs in the Red Sindhi, Kankrej, Gir, and Tharparkar assemblies were 134 Mb, 11.4 Mb, 7.5 Mb, and 6.2 Mb, respectively. Sahiwal and Red Sindhi assemblies demonstrated higher contiguity, with 90% of the genome assembled into 62 and 100 contigs, respectively. In contrast, Kankrej, Gir, and Tharparkar assemblies were less contiguous, featuring 5,472, 5,202, and 3,848 sequences, respectively, for L90. Finally, two pseudo-haplotypes were generated for each diploid genome to enable comparisons with the reference genome (**Supplementary Table S1)**.

**Table 1:**
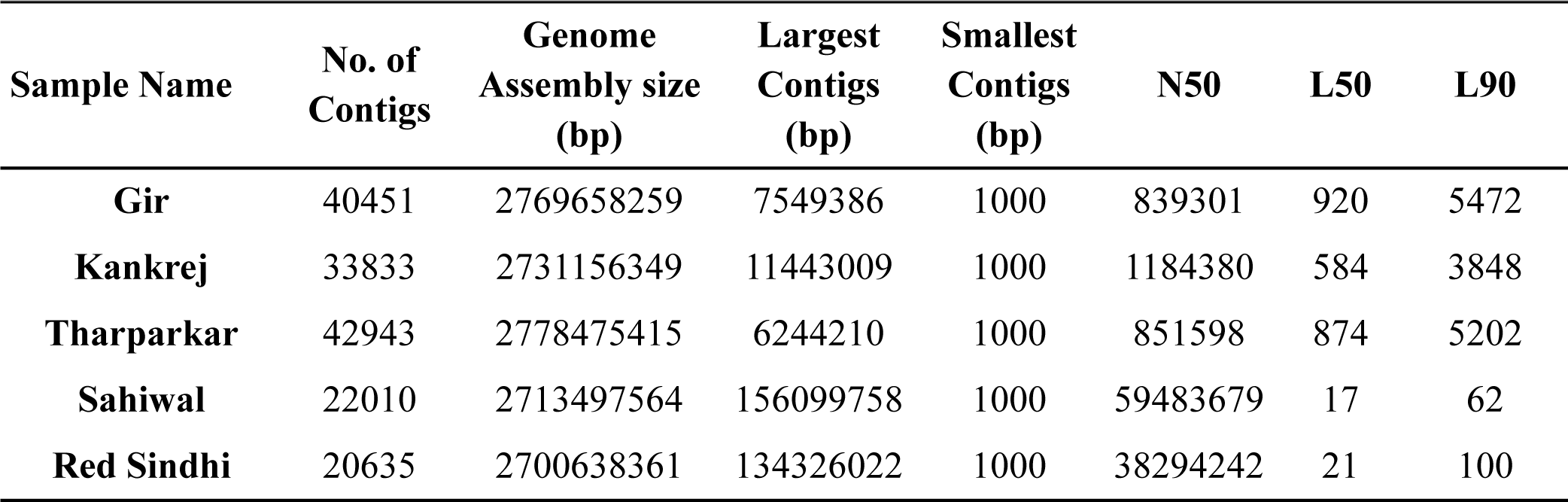
Data statistics of five *Bos indicus* de-novo genome assemblies.

### Iterative mapping based NUI discovery in *Bos indicus*

Upon completion, the NUI pipeline (**Fig. 1**) generated a FASTA file and a table representing the sequences of 9,270 non-redundant NUIs and their distribution in each sample, respectively (**Supplementary Table S2**). In further screening of these NUIs for contaminated sequences with stringent criteria, we identified and excluded 122 contaminated contigs, resulting in a clean set of 9,148 NUIs spanning 8 Mb. These NUIs ranged in size from 51 bp to 98.1 Kb. Among the five individuals, Gir had the fewest NUIs, totaling 1,972, while Red Sindhi had the highest number of NUIs, with 3,901 **(Supplementary Fig. S1)**. Notably, most NUIs were unique to each individual, with only 123 NUIs found to be common across all the individuals.

**Figure 1:**
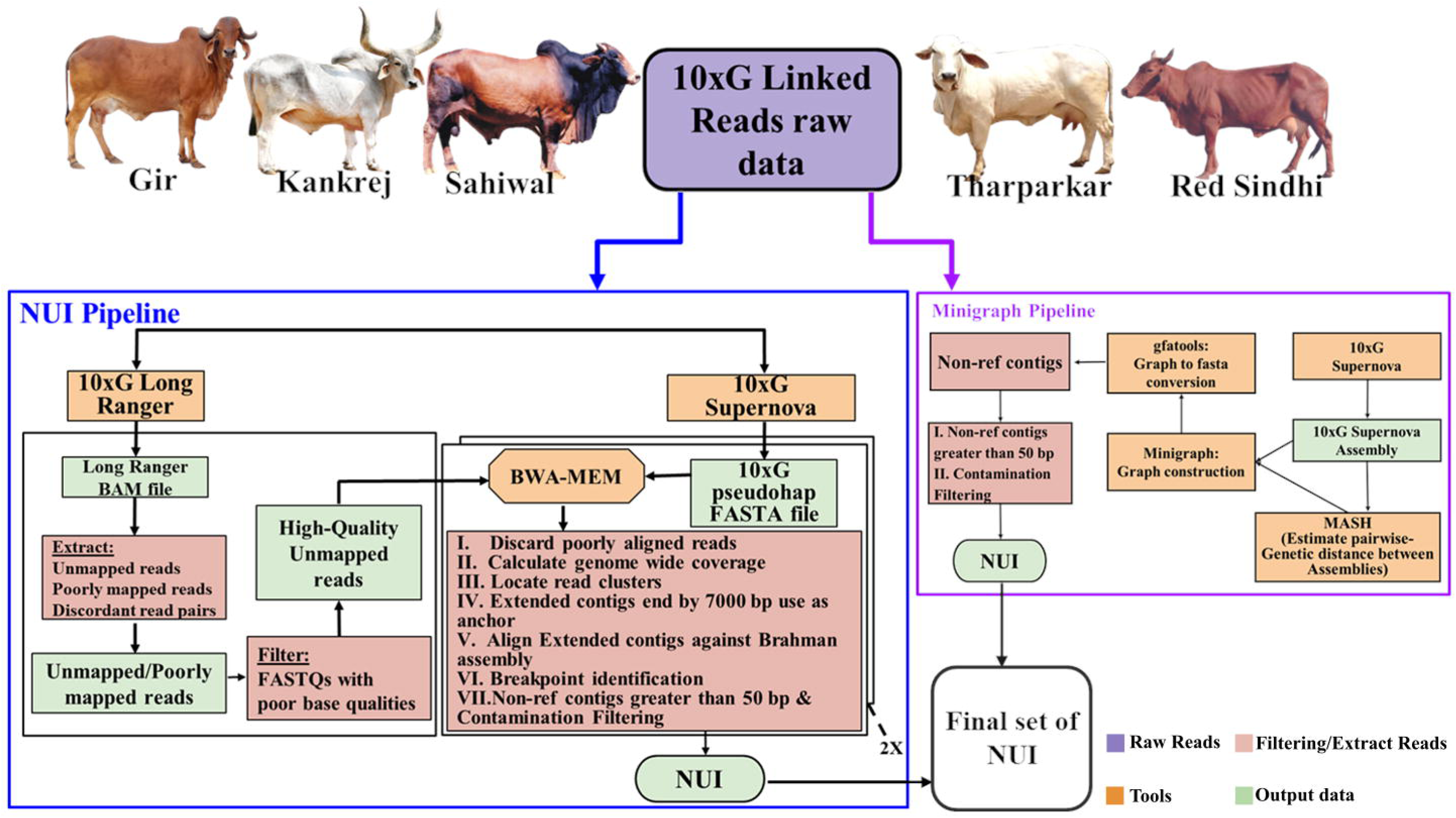
Identification of Non-reference Unique Insertions (NUIs) in *Bos Indicus*. The flowchart illustrates the systematic process for identifying the final set of NUIs. The diagram outlines the sequential steps involved in the selection and refinement of NUIs.

### Graph base NUI discovery in *Bos indicus*

A *Bos indicus* multi-assembly graph was constructed with the Brahman reference genome [32] as backbone. It contained a total of 2,728,215,813 bp across 153,597 nodes, connected by 218,354 edges. Specifically, there are 133,985 edges connecting two reference nodes, 83,089 edges connecting reference nodes to non-reference nodes, and finally, 1,280 edges connecting non-reference to non-reference nodes.

In the multi-assembly graph, a total of 2,709,654,022 bp consists of 111,178 reference nodes, which originated from the Brahman genome. This forms the backbone of the graph. With incremental integration of Kankrej, Gir, Sahiwal, Red Sindhi, and Tharparkar, the graph further expanded. Kankrej added 16,174 nodes with 7,369,711 bp, Gir contributed 9,306 nodes with 3,711,106 bp, Sahiwal accounted for 7,286 nodes with 3,124,476 bp, Red Sindhi introduced 5,536 nodes with 2,629,123 bp, and, lastly, Tharparkar added 4,117 nodes with 1,727,375 bp. Consequently, the multi-assembly graph comprises 42,419 non-reference nodes containing 18,561,791 bp **(Table 2)**. The non-reference nodes in the assembly graph are most abundant in Red Sindhi with 19,527 nodes featuring 8,667,230 bp. Gir features the least with 17,848 nodes encompassing 7,755,164 bp **(Table 3)**.

**Table 2:**
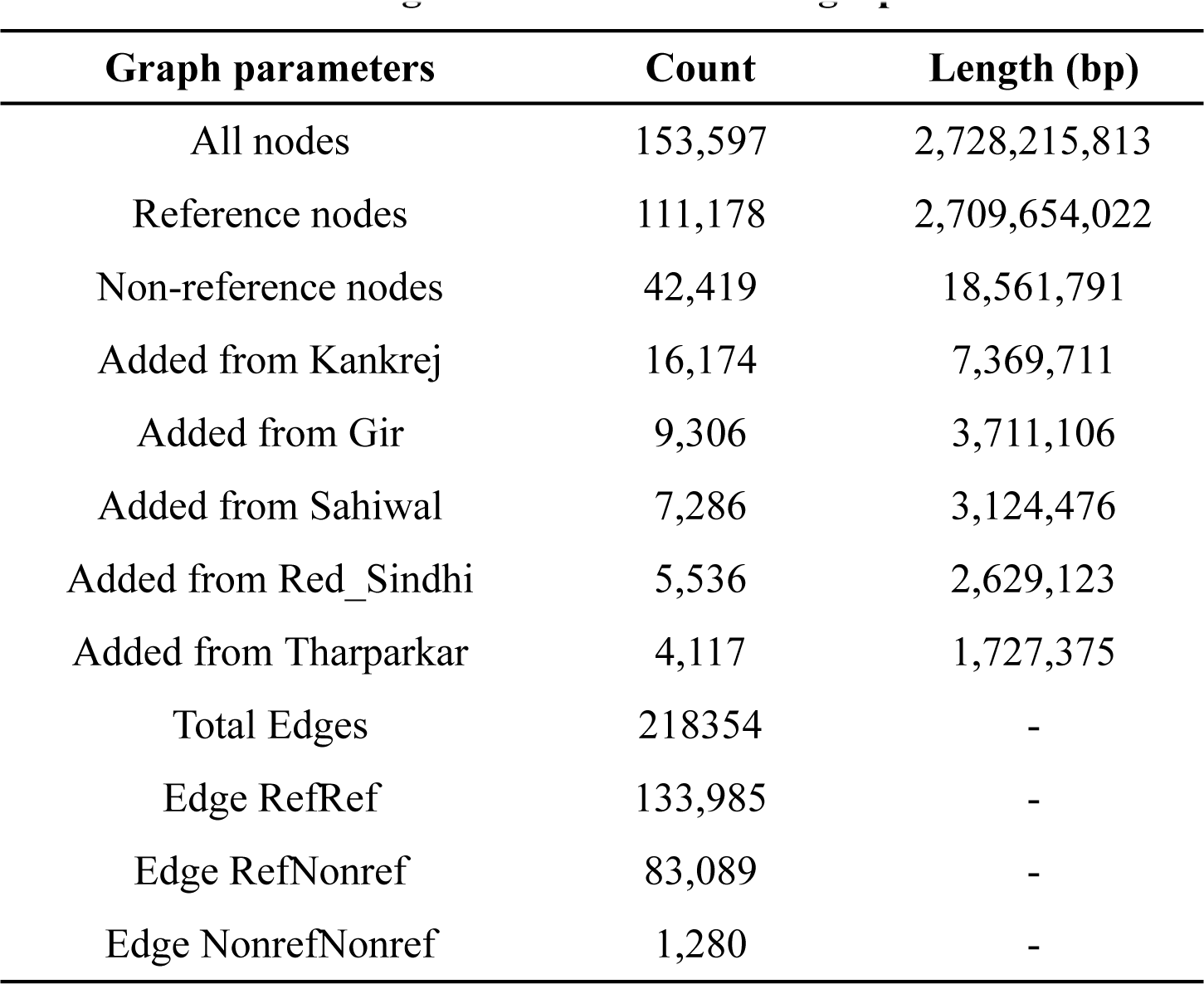
Nodes and edges statistics from minigraph.

**Table 3:** Nodes summary of five *Bos indicus* breeds from minigraph.

| **Non-ref sequences in Breed** | **Node count** | **Total sequence length (bp)** |
| --- | --- | --- |
| **Gir** | 17,848 | 7,755,164 |
| **Kankrej** | 18,252 | 8,506,737 |
| **Tharparkar** | 18,042 | 7,909,798 |
| **Sahiwal** | 19,195 | 8,477,068 |
| **Red Sindhi** | 19,527 | 8,667,230 |

The core genome of the multi-assembly graph, primarily the nodes shared by all assemblies, has a total count of 52,298 nodes featuring 2,399,340,052 bp. This constitutes 87.95% of the pangenome. Meanwhile, a total of 101,299 nodes, which constitute 12.05% of the pangenome, represent 328,875,761 bp and are considered flexible. This means these nodes do not appear in all assemblies. The flexible content of the genome is subdivided into two categories. Nodes that are present in at least two assemblies count for 76,322 with a total of 313,331,623 bp. Conversely, 24,977 nodes representing 15,544,138 bp are only found in a single assembly.

In the end, the non-reference nodes consisting of 18,561,791 bp were filtered to include only those greater than 50 bp. Subsequently, these selected nodes underwent a screening for contaminated sequences, leading to the identification of 29,477 clean nodes. These 29,477 clean nodes, spanning 17.73 Mb with the largest node of 98.1 kb, were retained as graph-based insertion sequences for further downstream analysis **(Supplementary Fig. S2)**.

### Selection of common set of NUI and comparison with published pangenomes

The NUI discovery pipeline and graph-based method ultimately provided clean datasets of 9,148 and 29,477 insertions respectively. A comprehensive comparison of these two datasets revealed 6,844 NUIs within the graph-based insertion sequences, establishing them as a confident and final set of NUIs spanning 7.57 Mb. All these NUIs are breakpoint-resolved and precisely located the chromosomes of the Brahman reference genome **(Fig. 2A;Fig. 2B)**. This invaluable breakpoint information facilitates downstream analysis in the population.

**Figure 2:**
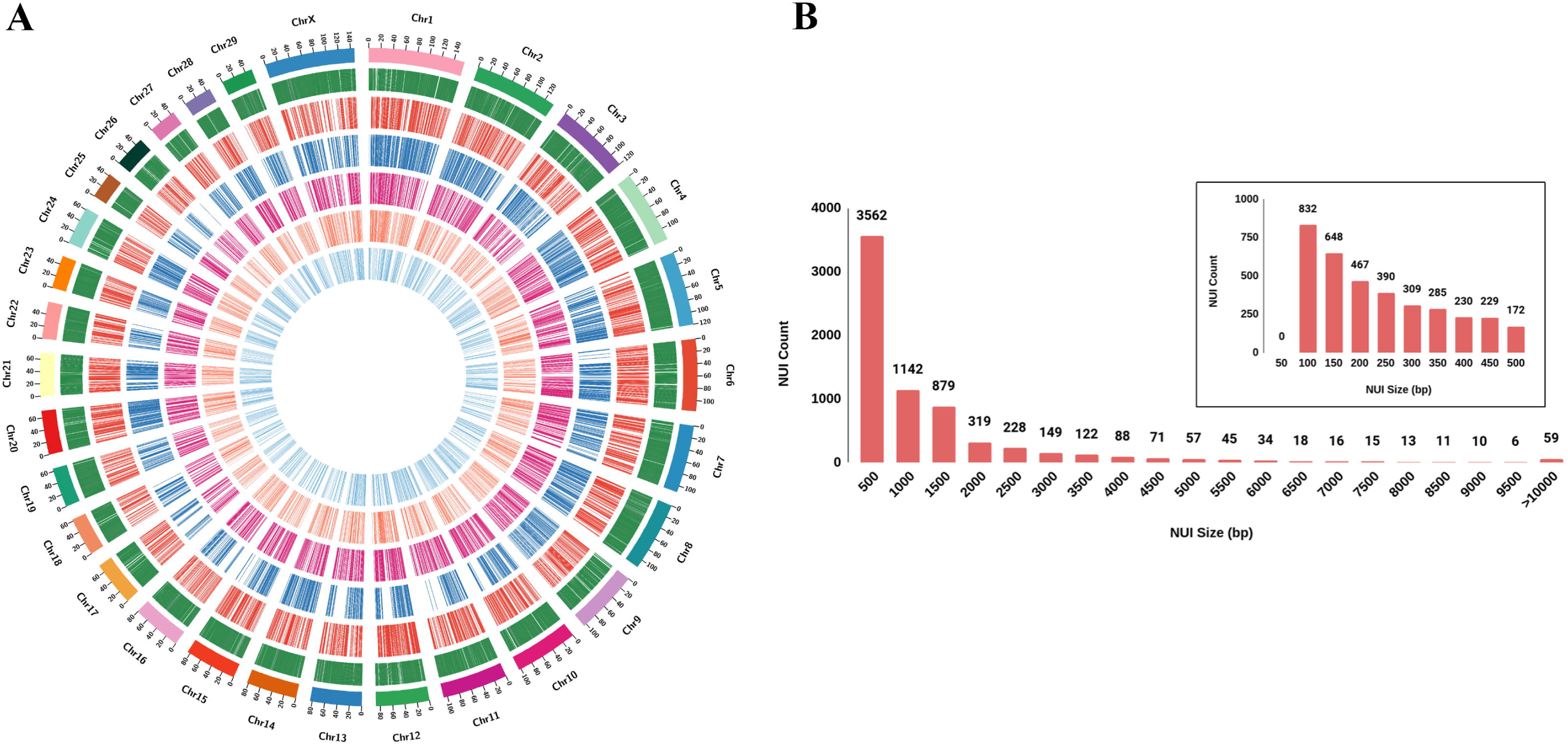
Overview of Non-Reference Unique Insertion (NUI) final set and their distribution. **(A)** Circos plot of *Bos indicus* pangenome, From the outer to inner track: Chromosome, Gene track, Red Sindhi, Sahiwal, Tharparkar, Kankrej, Gir **(B)**Size distribution of NUIs with bin size of 500 bp and 50 bp in zoomed area.

Subsequently, these NUIs were compared with two published cattle pangenomes. When compared with the comprehensive cattle pangenome constructed by Zhou et al. [12], consisting of 22,324 contigs compiled from 898 cattle of 57 breeds, only 258 NUIs (∼4%) spanning 76.92 Kb (∼1%), were found to be in common. This may be attributed to the fact that the dataset primarily included European and African cattle breeds, with relatively limited inclusion of Indian breeds. Specifically, only the Gir breed from our study was featured, alongside other *Bos indicus* breeds like Brahman and Nellore.

In contrast, the comparison with the recently published pangenome of *Bos indicus* cattle [19], developed from genome assemblies of 10 Chinese indicine breeds, demonstrated a more substantial overlap. In fact, when the NUIs identified in this study were compared with 74,907 Chinese pangenome sequences spanning 124.4 Mb, it was observed that 2.80 Mb (∼37%), consisting of 3,712 NUIs (∼54%), were in common. Smaller contigs were more likely to overlap with 58% of contigs less than 1Kb matching (**Supplementary Table S3**)

### Screening for NUIs in the Population and Identification of BICIs

The evaluation of NUIs within a population of the same cattle breeds is vital for understanding their distribution, variability, and validation. In this study, we conducted genotyping for a total of 6,844 NUIs across 98 individuals, encompassing 20 Gir, 19 Kankrej, 20 Tharparkar, 20 Sahiwal, and 19 Red Sindhi cattle. This genotyping effort revealed the presence of both rare and common NUIs within the population. In fact, 2,789 NUIs were not called in any samples whereas 1,091 NUIs were present across all samples. To focus our analysis on the common NUIs and exclude the rare ones, we applied a stringent filtering criterion based on a Minor Allele Frequency (MAF) threshold of < 5%. As a result, we established a final NUIs call set comprising 2,312 NUIs, collectively spanning ∼2 Mb of genomic sequence **referred to as BICIs**. The BICIs were distributed across all the chromosomes (**Fig. 3A**). Among these, the largest BICI measured 47.6 Kb, whereas the smallest were as short as 52 bp. It’s noteworthy that 25% of BICIs (578) were equal to or less than 130 bp in length, and 50% of BICIs (1,156) were 286 bp or shorter (**Fig. 3B**). This refined BICIs dataset represents the common NUIs in the population and lays the foundation for all subsequent analyses in this study.

**Figure 3:**
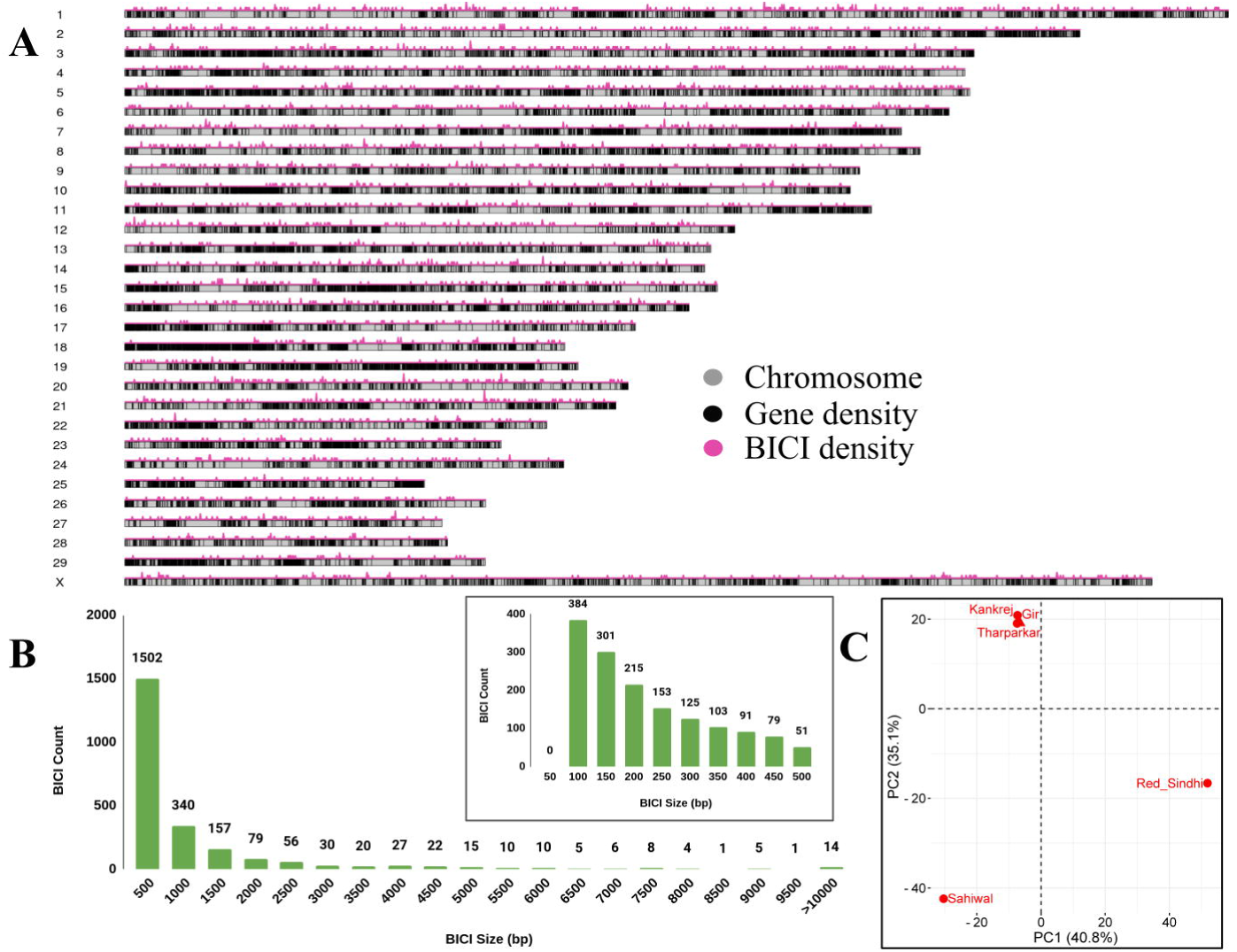
Overview of *Bos Indicus* Common Insertions (BICIs) and their distribution patterns. **(A)** This ideogram depicts BICIs occurrences across Brahman genome chromosomes. Pink histograms above each chromosome illustrate BICI density using a 100 kb window size. The gray line within chromosomes represents gene distribution. **(B)** The plot illustrates the size distribution of BICIs with a bin size of 500 bp, providing an overview and 50 bp in the zoomed area. **(C)** The first two principal components are based on the BICI occurrence matrix.

### Principal Component Analysis and hierarchical clustering of BICIs

The distribution of the 2,312 BICIs was assessed across the five genome assemblies (**Supplementary Table S4**). Notably, a large number of unique BICIs were identified in Red Sindhi (638 BICIs), followed by Sahiwal (591 BICIs), while the fewest BICIs were found in Tharparkar assembly (97 BICIs). Additionally, 16 BICIs were present in all five assemblies.

Principal Component Analysis (PCA) applied to these 2,312 BICIs across the five distinct assemblies effectively differentiated the Gir, Kankrej, and Tharparkar breeds from the Sahiwal and Red Sindhi breeds. PC1 explained 40.8% of the variation, while PC2 accounted for 35.1%. A PCA1 vs. PCA2 plot in **(Fig. 3C)** provided a clear separation of animals from different geographical regions, underscoring the utility of BICIs in discerning breed differences.

The genotyping results, depicting the presence and absence of BICIs in the population, underwent hierarchical clustering analysis. This analysis resulted in the formation of major clusters representing different breeds. Notably, Red Sindhi and Gir each formed distinct clusters, as did Kankrej and Sahiwal. Tharparkar individuals were an exception and could not be grouped into a single cluster. In fact, they clustered into two groups, with the larger group having one Sahiwal individual and four Kankrej individuals that did not group with their own cluster. The resulting heatmap visually illustrates the distribution of 2312 BICIs in the population (**Supplementary Fig. S3**).

### Identifying the major transposable element in BICIs

In our investigation of repeats and the composition of TEs within BICIs, we discovered that approximately 63.21% of the bases within the BICI call set contained interspersed repeats (**Supplementary Table S5**). This proportion is notably higher than the total interspersed repeats found in the entire genome of the Brahman reference sequence (46.71%). Further analysis revealed that 11.32% of the BICIs were short interspersed nuclear elements (SINEs), while long interspersed nuclear elements (LINEs) constituted 43.87% of the overall BICI dataset. This contrasts with the genome-wide repeat content of SINEs and LINEs in the Brahman reference genome, which accounted for 11.75% and 27.94%, respectively. It is evident that LINEs are significantly enriched within the BICI dataset, highlighting the prevalence of this particular repetitive element in these insertions. We further explored the distribution of major TE types within BICIs based on their sizes. Across all size categories, LINEs were the most prevalent TE within BICIs. The proportion of LINEs increased from 40.91% in BICIs under 200 bp category to a substantial 69% in longer sequences exceeding 1000 bp. Conversely, BICIs under 200 bp were predominantly composed of non-repetitive unique sequences, while the occurrence of unique sequences declined as BICI size increased. Long terminal repeats (LTRs) and DNA transposons were observed at lower frequencies within all BICI size ranges, suggesting their relatively limited representation within the BICI dataset regardless of BICI **(Fig. 4A)**. These findings underscore the dominance of LINEs and the influence of BICI size on repeat content.

**Figure 4:**
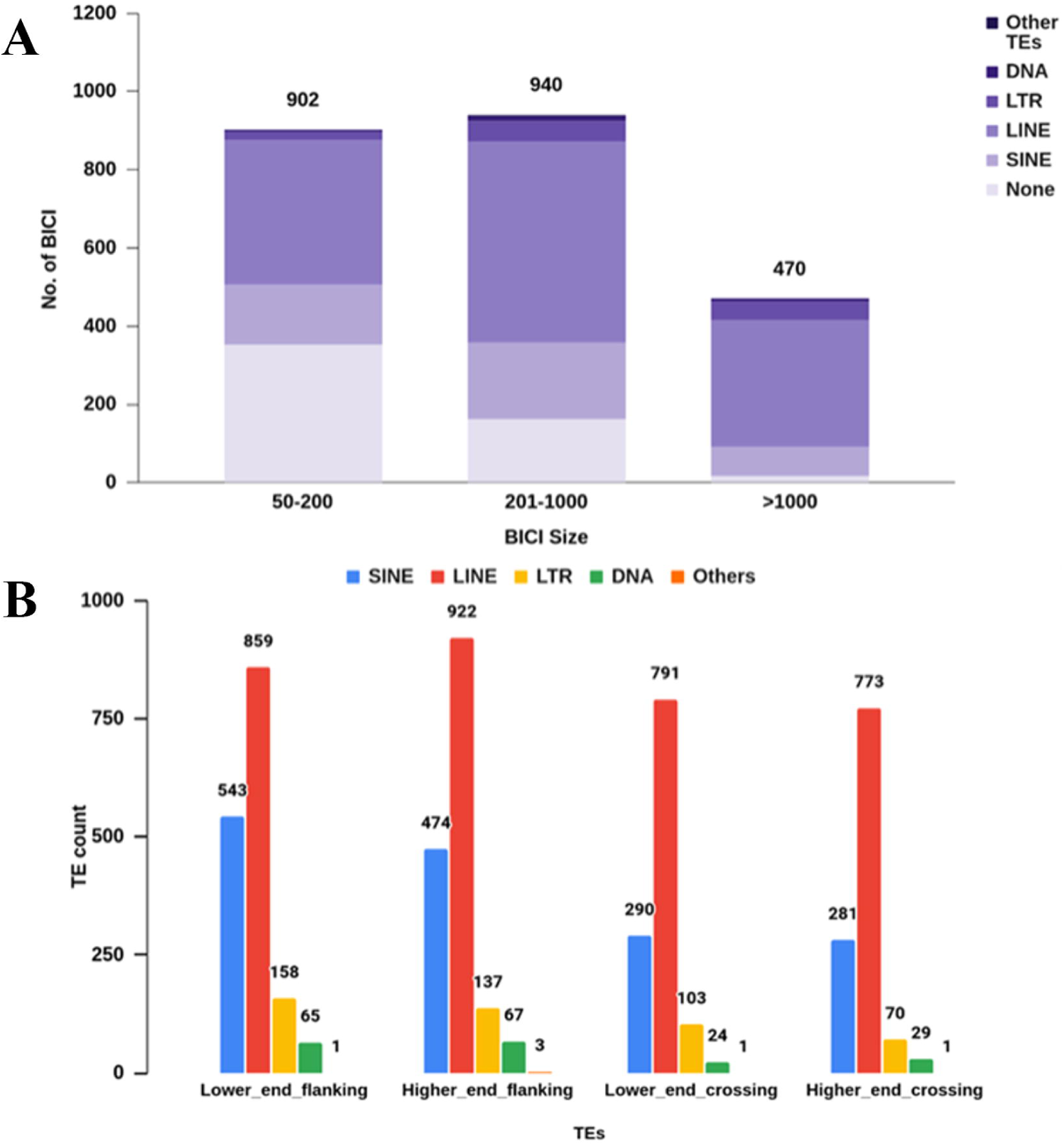
Distribution of Transposable elements on *Bos Indicus* Common Insertions (BICIs). **(A)** Stacked bars represent the total number of BICIs split by three different size ranges. The major transposable elements (TEs) are categorized as SINE (Short Interspersed Nuclear Element), LTR (Long Terminal Repeat), LINE (Long Interspersed Nuclear Element), DNA (DNA Transposon), and NONE (No Interspersed Repeat Detected). “Other TEs” encompasses various minor classes.

We additionally characterized the repetitive sequences in the flanking regions of BICIs and found that approximately 70% of the BICIs were flanked by a transposable element (TE) on at least one end. This underscores the association of BICIs with TE sequences in their vicinity. Furthermore, we found that around 50% of BICI breakpoint crossing over sequences were encompassed by TEs, emphasizing the substantial impact of TEs on BICI breakpoints **(Fig. 4B)**. This dual observation underscores the intricate interplay between BICIs and TEs in shaping the cattle genome, shedding light on their interdependencies and potential functional roles.

### Gene coding and Transcriptional potential of BICIs

The genomic distribution and classification of BICIs were examined by pinpointing their breakpoints within the Brahman reference genome annotation. Out of the 2,312 BICIs in our dataset, 926 BICIs were positioned within genic regions, while the remaining 1,386 BICIs were situated outside of annotated genes (**Supplementary Table S6**). A detailed analysis of these BICIs revealed that 26 of them were in exons, highlighting their potential to influence protein-coding sequences. The presence of BICIs was associated with a total of 754 genes, demonstrating the diversity of genic regions impacted by these insertions. Remarkably, 121 genes within this group contained two or more BICIs within their span, underscoring the presence of multiple BICIs within specific genomic loci. The majority of genes linked with BICIs were classified as protein-coding genes, strengthening the notion that BICIs may have a functional impact on protein-coding sequences. Additionally, BICIs were found within 54 long non-coding RNA (lncRNA), while few BICIs were associated with 18 pseudogenes **(Table 4)**. These findings provide a comprehensive perspective on the distribution and potential implications of BICIs in various genomic contexts, including protein-coding regions, non-coding sequences, and pseudogenes.

**Table 4:** Statistics of BICIs annotated in gene. BICIs in Different Genomic Features

| **BICIs in Different Genomic Features** |  |
| --- | --- |
| BICIs in Gene | 926 |
| BICIs in Exon | 26 |
| BICIs in Intron | 900 |
| BICIs Intergenic | 1,386 |
| **Non-Redundant Gene set Associated with BICIs:** |  |
| Total Non-Redundant Genes | 754 |
| Protein coding genes | 682 |
| lncRNA | 54 |
| Pseudogene | 18 |

While the number of BICIs situated within exons is vanishingly small, approximately 0.02%, it remains plausible that some may represent previously unseen exons or regulatory elements with transcriptional potential. To address this, we employed both evidence-based and ab-initio methods to assess the transcriptional potential of the BICI dataset. Utilizing high-quality RNASeq data from 47 in-house samples derived from peripheral blood mononuclear cells (PBMCs) for evidence-based annotation, we identified 1,631 transcripts originating from 1,395 BICIs. Among these, 662 transcripts from 602 BICIs were located within genic regions, while 969 transcripts were associated with 793 BICIs located outside the genic regions of the reference genome (**Fig. 5A**; **Supplementary Table S7**). Furthermore, our ab-initio analysis revealed the transcription potential of 148 genes from 124 BICIs. Of these, 137 were supported by transcripts, with 37 genes from 29 BICIs situated within genic regions and 100 transcripts from 85 BICIs having breakpoints outside the genic regions (**Supplementary Table S8**). Of the 11 genes associated with 10 BICIs that lacked transcript support, 7 BICIs were situated in non-genic regions and 3 BICIs were located within genic regions of the Brahman reference genome.

**Figure 5:**
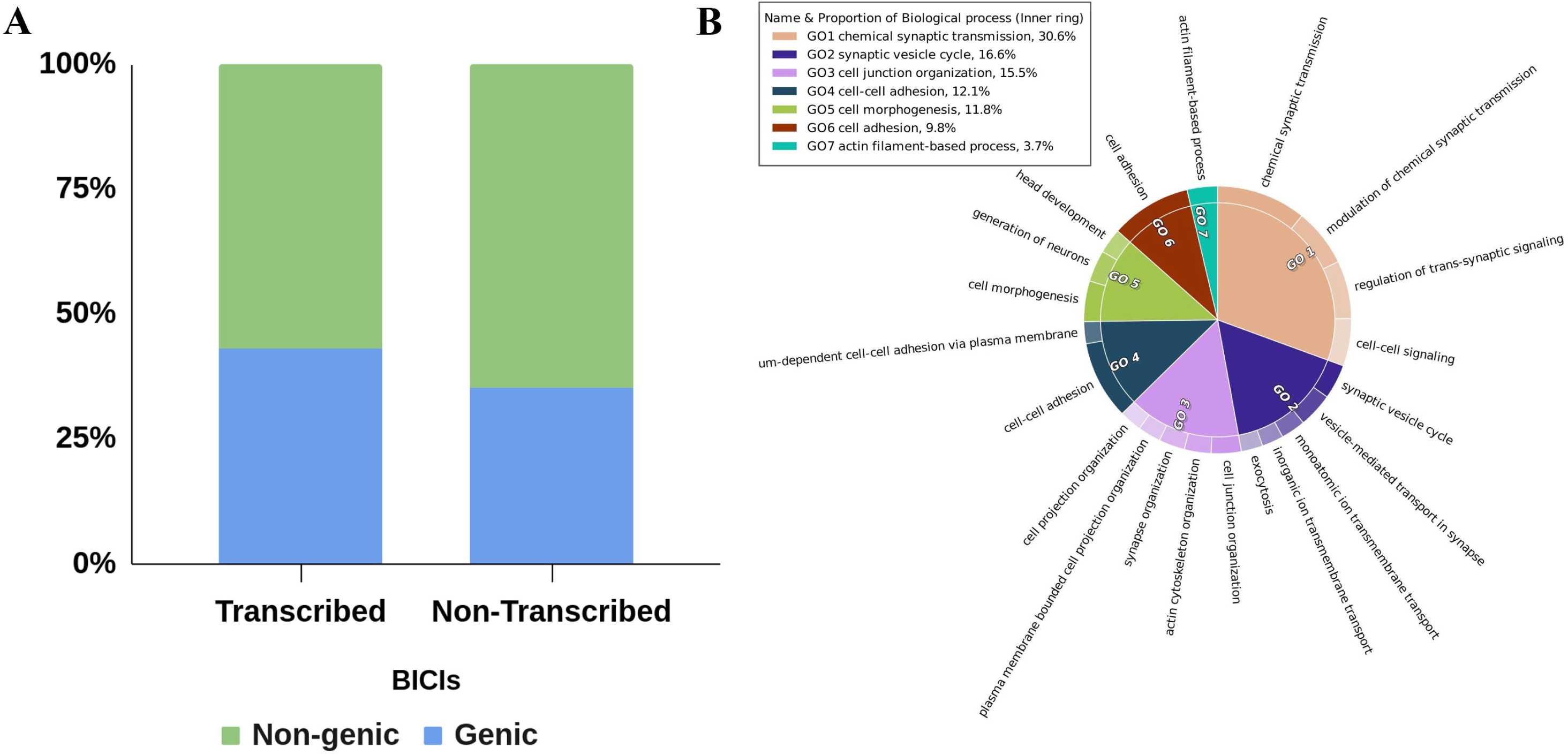
Characterization of *Bos Indicus* Common Insertions (BICIs) in the transcriptome. **(A)** Stacked bars depict the distribution of BICIs across genic and non-genic regions of the genome. **(B)**Gene Ontology (GO) annotation analysis for the biological processes associated with BICIs.

### Impact of BICIs in functional genome

The functional analysis of protein coding genes with BICI unveiled enrichment in genes predicted to be associated with key biological processes. Notable enriched terms included chemical synaptic transmission, cell junction organization, cell-cell adhesion, and cell morphogenesis (**Fig. 5B**). Simultaneously, the examination of major cellular components highlighted involvement in the synapse, plasma membrane region, and monatomic ion channel complex (**Supplementary Fig. S4A**). Furthermore, the Gene Ontology (GO) enrichment analysis for molecular functions indicated enrichment in ion channel activity, glutamate receptor activity, cell adhesion-mediated activity, and 3’,5’-cyclic-AMP phosphodiesterase activity (**Supplementary Fig. S4B).**

QTL analysis was conducted on 682 protein coding genes with BICIs, revealing approximately 3,368 QTLs associated with 539 genes. Five genes lacked a corresponding location in the ARS-UCD1.2 reference genome [22] and 138 genes did not exhibit enrichment with the cattle QTL database. As expected, several QTLs were associated with individual genes. These QTLs were categorized into six main classes: milk production, reproduction, exterior traits, health, meat production, and carcass traits. Notably, milk production traits (28.7%) were the most enriched category, followed by exterior traits (17.5%) (**Fig. 6A**). Enrichment analysis further revealed that top QTLs predominantly influenced milk traits (milk yield, milk fat yield, milk fat percentage, milk protein yield, and milk protein percentage) and production traits (body weight, and body weight gain) (**Fig. 6B**). The integration of functional analysis with QTL mapping revealed a strong link between genes with BICIs and economically important traits in *Bos indicus* cattle. Our analysis of common genes within enriched GO terms further revealed overlap with enriched QTLs. Notably, genes like CTNNA3 and CTNNA2, enriched in GO terms for ’cell adhesion’ and ’cell migration’ respectively, reside within the milk QTL region. Similarly, the ROBO1 gene, enriched in GO terms related to ’cell adhesion’ and ’migration,’ is also present in the reproduction trait QTL. Furthermore, major genes like ITPR1 and ITPR2, enriched in the GO terms related to ’cell morphogenesis,’ ’calcium ion transmembrane transport,’ and ’response to hypoxia,’ are identified within the body weight QTL region.

**Figure 6:**
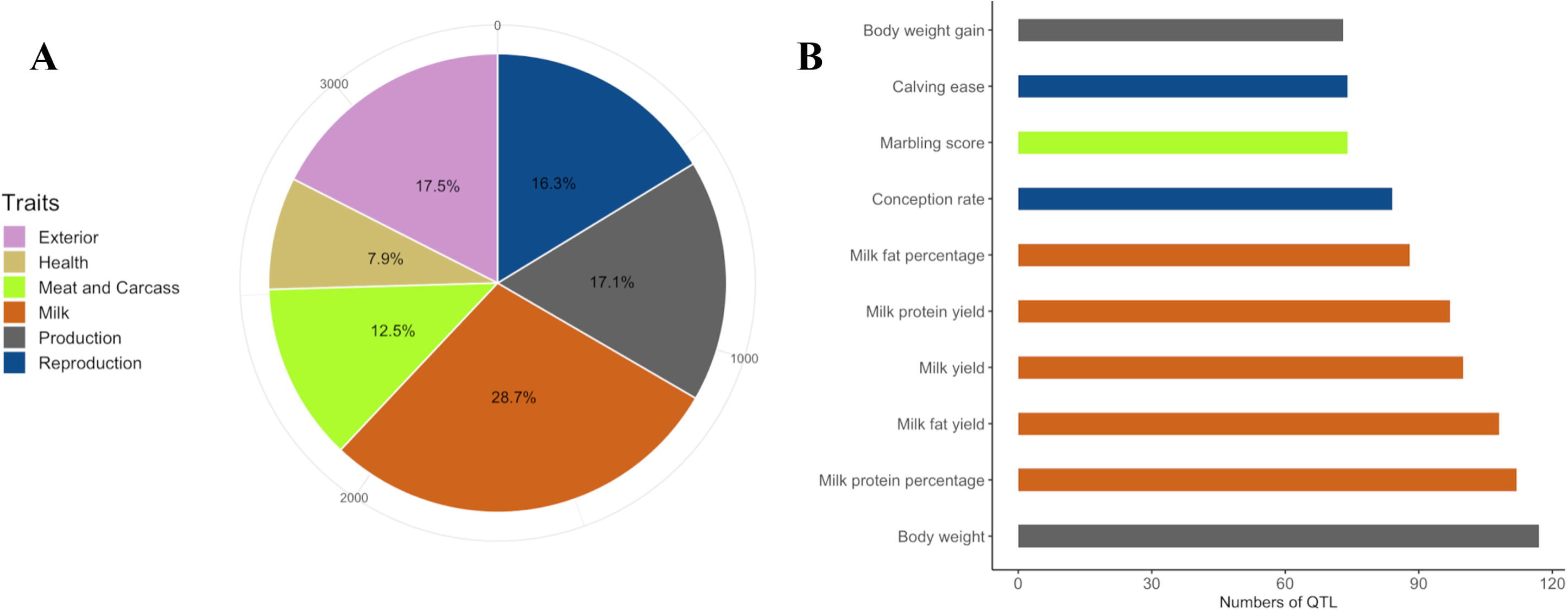
Trait enrichment analysis of BICIs in known cattle QTL regions. **(A)** Percentage of QTL type (pie chart) associated with BICIs. **(B)** Top ten enriched QTL traits (bar plots) associated with BICIs.

### Evolutionary analysis of the BICIs

To elucidate the evolutionary origins of the BICIs, we conducted alignment analyses with sister species of the Bos genus and other evolutionarily related species within the Bovidae family. Specifically, we aligned BICIs to five Bos species, which included *Bos taurus* (taurine cattle), *Bos gaurus* (gaur), *Bos frontalis* (gayal), *Bos grunniens* (domestic yak), and *Bos mutus* (wild yak). Out of the total 2,312 BICIs, a substantial proportion, 1,625 (70.28%), was identified across the Bos sister species. Each sister species exhibited alignment with over 40% of the BICIs. Specifically, *Bos taurus* aligned with 1,002 BICIs (43.4%), *Bos gaurus* with 1,172 BICIs (50.7%), *Bos frontalis* with 950 BICIs (41.1%), *Bos grunniens* with 1,152 BICIs (49.8%), and *Bos mutus* with 1,156 BICIs (50%) **(Fig. 7A)**. Notably, most BICIs were not exclusive to a single sister species but were shared across multiple sister species. Specifically, 467 (20.2%) BICIs were common to all five sister species, while 415 BICIs were shared with at least four sister species. Additionally, there were a few BICIs that were unique to specific sister species, including *Bos taurus* (218), *Bos gaurus* (51), *Bos frontalis* (28), *Bos grunniens* (13), and *Bos mutus* (3). This distribution of unique BICIs aligns with the phylogenetic tree and genetic introgression observed in the Bos genus as previously described by Wu et al. [34].

**Figure 7:**
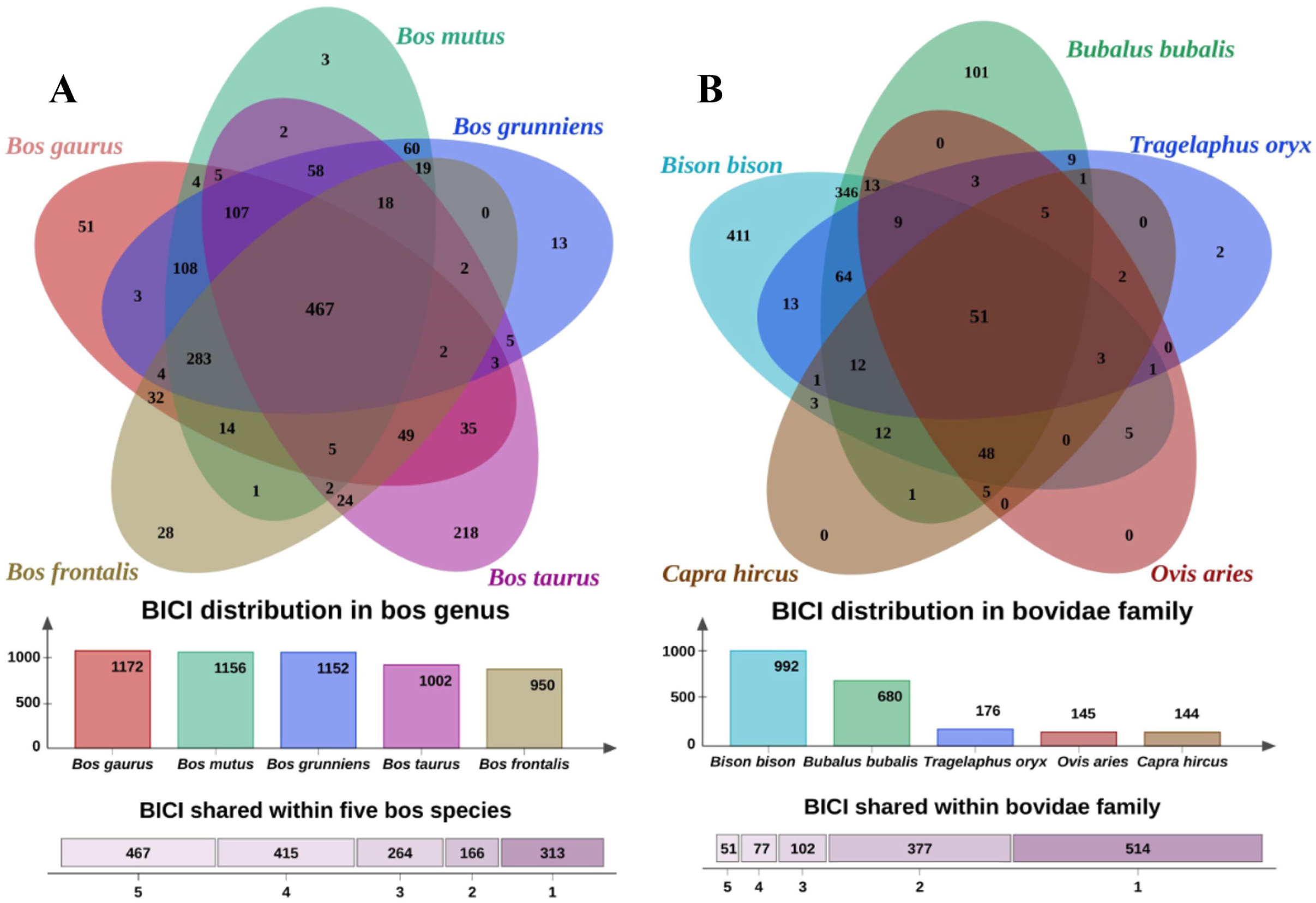
Evolutionary analysis of *Bos Indicus* Common Insertions (BICI). **(A)** Venn diagram illustrating the number of *Bos Indicus* Common Insertions (BICIs) shared within the Bos genus. The overlapping regions reveal the extent of shared BICIs among species within the Bos genus. **(B)**Venn diagram showcasing the number of *Bos Indicus* Common Insertions (BICIs) shared within the Bovidae family. The intersecting areas depict the shared BICIs among species within the broader Bovidae family.

Similarly, when BICIs were aligned with more distantly related species within the Bovidae family, such as American bison (*Bison bison*), eland (*Tragelaphus oryx*), buffalo (*Bubalus bubalis*), sheep (*Ovis aries*), and goat (*Capra hircus*), only 1,121 BICIs aligned to any of these genomes, which accounted for less than 50% of the BICIs. Most BICIs matched with the American bison (992), followed by buffalo (680), eland (176), sheep (145), and goat (144) **(Fig. 7B)**. Regarding the distribution of matched BICIs, the lowest number of BICIs (51) were shared across all five genomes, followed by 77 shared across four genomes, 102 shared across three genomes, 377 shared across two genomes, and 514 were unique to specific species. Particularly noteworthy is the prevalence of unique BICIs in the bison genome (411), followed by buffalo (101) and eland (2). Goat and sheep did not exhibit any unique BICIs within their genomes. This distinctive BICI distribution across these species highlights their evolutionary divergence and lineage-specific enrichment of BICIs.

## Discussion

The present study is dedicated to uncovering the presence and characteristics of NUIs, representing a substantial segment of the cattle pangenome that is absent in the *Bos indicus* reference genome. To unveil these NUIs, the genomes of five desi cattle breeds, Gir, Tharparkar, Kankrej, Sahiwal, and Red Sindhi, were sequenced utilizing 10X Chromium technology [33]. The length of these genome assemblies were comparable to the published Brahman reference genome and other cattle genomes assembled using long-reads [19,22,32,35]. However, it was evident that the contiguity of scaffolds in the assemblies varied. This variance can be attributed to the intricacies of linked-read library preparations [33,36]. Notably, the assembly size was found to be contingent upon the molecular size selected for linked-read preparation.

The evolution of methods for comparing genomes with reference sequences to identify missing segments has been noteworthy. Initially, direct alignment-based methods were employed for NUI detection; however, graph-based methods have gained prominence for their purported comprehensiveness. In our study, we utilized a pipeline specifically tailored for linked reads, developed by Wong et al. [29] for diploid species and applied to humans. Concurrently, we constructed the pangenome using a graph-based approach, revealing a more extensive set of NUIs spanning 17.7 Mb compared to the 8 Mb identified through the NUI discovery pipeline. Although the pipeline relies on the indirect comparison of five genomes by aligning short reads, it is noteworthy that, out of the 8 Mb identified, 7.5 Mb was also corroborated by the graph-based approach. This observation underscores the stringency of the pipeline, ensuring the identification of a confident set of NUIs. The advantage of the pipeline lies in its ability to precisely report all NUIs for which breakpoints in the reference genome are identified. It accurately pinpoints the location of these NUIs, enhancing confidence in their identification. While alignment methods have traditionally dominated NUI detection, the prevalent trend in pangenome approaches involves the utilization of graph-based methods. This study contributes to the growing body of evidence supporting the effectiveness of both alignment and graph-based methods, emphasizing their complementary roles in constructing a comprehensive understanding of the cattle pangenome.

It is imperative to underscore the distinctive focus of our study, which centers on the Brahman reference genome and marks the pioneering effort in constructing a pangenome specific to desi cattle of the subspecies *Bos indicus*. This exclusive emphasis on a specific subspecies serves as a departure from other recent cattle studies, such as those conducted by Zhou et al. [12] and Crysnanto et al. [23], which reported larger pangenome sizes of 83 Mb and 70 Mb respectively. The contrasting sizes reported in these studies can be attributed to their broader inclusion of diverse breeds, encompassing not only different subspecies but also wild relatives. Notably, the inclusion of a greater number of distant breeds tends to augment the overall pangenome size by capturing a more extensive array of genetic variations.

NUIs identified in the study were subjected to a comprehensive characterization based on the presence and absence variations in a population of 5 desi breeds. The focus was on identifying NUIs that displayed frequent and common occurrences, as such variants often play a more substantial role in influencing traits and diseases. Common variants not only exhibit more reliable associations in genetic studies but also possess the potential for broader implications, given their prevalence in diverse populations [37–40]. Consequently, 2,312 NUIs, spanning approximately 2 Mb, were selected as common NUIs and referred to as BICIs, with a minimum allele frequency (MAF) > 0.05. It is noteworthy that the majority of the identified NUIs in our study were not classified as common, a pattern that aligns with findings in the African pangenome, where a significant portion of variants was reported as private to individuals. Additionally, in another study focusing on the *Bos indicus* pangenome using short reads (unpublished), a substantial number of NUIs were identified as private to specific samples. Importantly, this filtering process also led to the exclusion of NUIs that were present across all samples. These are the NUIs that are only absent in the Brahman genome, suggesting that the Brahman, an indicus breed from the USA, exhibits genetic distinctions from other Indian indicus breeds. Furthermore, this observation raises the possibility that some NUIs may be present in the Brahman genome but not captured during the assembly process.

The identification of a higher number of NUIs in Red Sindhi and Sahiwal assemblies underscores the significance of a contiguous genome for accurate NUI detection. The presence of BICIs within assemblies facilitates a clear differentiation between breeds, as demonstrated through PCA where animals were separated based on their geographical origin. Genotyping these BICIs in the population revealed a clustering of Red Sindhi, Gir, and Sahiwal, pointing to a genetic distinctiveness among these breeds. However, Tharparkar and Kankrej exhibited a more mixed clustering pattern, suggesting potential genetic similarities or variations between these breeds, which is noteworthy given their geographical proximity. This proximity might contribute to shared genetic traits or interbreed variations, influencing their genomic profiles. The unique clustering pattern observed not only emphasizes the utility of BICIs as markers for capturing and characterizing genetic diversity between cattle breeds but also highlights their practical application for breed differentiation. The regional clustering aligns with findings from other studies employing different marker systems, providing further validation for the consistency of breed-specific genomic signatures.

The exploration into repeats and TEs within BICIs reveals intriguing insights into the cattle genome. In contrast to humans, where short interspersed nuclear elements (SINEs) dominate non-reference insertions [29], cattle exhibit a prevalence of long interspersed nuclear elements (LINEs) in BICIs. The presence of TEs in the flanking regions of BICIs mirrors human patterns [29], with approximately 70% of BICIs flanked by TEs at least on one end. This shared dynamic emphasizes the role of TEs in both BICIs and their adjacent sequences. The significance of the enrichment of TEs in non-reference insertions has been noted in various reports [41–43], indicating their potential role in BICIs genesis. The notable enrichment of LINEs in cattle prompts further exploration of their functional implications in shaping the cattle genome, building on previous suggestions regarding the significance of TEs in BICIs genesis.

The exploration of transcriptional potential through both ab-initio [44,45] and evidence-based methods [46,47] establishes that BICIs exhibit transcriptional activity. The predominant annotation of BICIs with protein-coding genes underscores their potential functional roles. Concurrently, the annotation of BICIs within long non-coding RNA (lncRNA) genes signifies their possible regulatory roles in the genome [48,49]. The presence of BICIs associated with pseudogenes further hints at their diverse functional repertoire [50]. Further, the identification of BICIs within intronic regions, displaying transcriptional potential, suggests the possibility of contributing additional sequences to existing transcripts. The prevalence of BICIs within genes is a common phenomenon, and the observed variation in their presence may contribute to alterations in gene length within a population [16]. In summary, the transcriptional profiling of BICIs provides compelling evidence for their functional significance, with potential roles in both coding and non-coding genomic elements. The diverse genomic locations and associations with different gene types underscore the intricate and multifaceted impact of BICIs on the regulatory landscape of the genome.

Genes found in the top enriched GO Biological Process (BP) categories also exhibit significant enrichment within major Quantitative Trait Locus (QTL) regions, supporting a functional link between biological processes and genetic variation. CTNNA3 plays a crucial role in the formation of cell-cell adhesion complexes, potentially influencing milk production, milk protein content percentage, milk protein yield, milk fat content percentage, and milk fat yield through its role in mammary gland development [51]. CTNNA2, found to be positively selected in the Gir dairy breed [52], plays a role in the development of the nervous system and has also shown association with climate adaptation in Mediterranean cattle [53]. The ROBO1 gene is enriched in GO terms related to ’cell adhesion’ and ’migration’, potentially impacting fertility and ovarian health [54]. ITPR1 has been reported to be associated with environmental high-altitude adaptation in the yak [55], potentially influencing body weight regulation. Additionally, ITPR2 has been linked to heat stress response in US Holsteins [56]. These findings suggest a potential role for these genes in mediating various phenotypic traits through their involvement in crucial biological processes.

The comprehensive alignment analyses aimed at unraveling the evolutionary origins of BICIs provided valuable insights into their distribution among sister species within the Bos genus and other evolutionarily related species in the Bovidae family. Notably, a substantial proportion (70.28%) of BICIs was identified across the Bos sister species, including *Bos taurus* (exotic cattle), *Bos gaurus* (Gaur), *Bos frontalis* (Gayal), *Bos grunniens* (Domestic Yak), and *Bos mutus* (wild Yak). Interestingly, the observation that the number of BICIs shared solely with *Bos taurus* was not the highest of challenges expectations. The lower total number of BICIs found in *Bos taurus* compared to other wild relatives suggests that *Bos taurus* may have undergone artificial selection [57], resulting in the loss of many BICIs. This is particularly evident in the subset of 283 BICIs shared with all other sister species. Further examination of distantly related species within the Bovidae family, such as American bison, eland, buffalo, sheep, and goat, revealed a varied distribution of matched BICIs, aligning with the inferred evolutionary divergence times based on phylogenetics [34]. This trend of shared and unique BICIs across species is reminiscent of findings in human genomics when NUIs of humans were compared with other species like chimpanzee, gorilla, orangutan, and bonobo [29]. Such comparative analyses not only highlight the evolutionary dynamics within the Bovidae family but also draw parallels with similar studies in different species, providing a broader understanding of the genomic changes accompanying evolutionary divergence.

## Conclusions

This study addresses the crucial need to explore and comprehend the genomic diversity within the *Bos indicus* population, with a specific focus on dairy breeds in India. The construction of the *Bos indicus* pangenome, employing alignment and graph-based methods, revealed significant differences in size, emphasizing the importance of diverse approaches. A robust set of NUIs spanning 7.8 Mbs which are common to both methods, defines the pangenome of Indian *Bos indicus* breeds. Comparative analyses showcased distinctions with other pangenomes, highlighting the unique genomic landscape of these dairy breeds. The identification of BICIs, particularly within protein-coding genes enriched for specific functions, provided insights into potential roles in various traits related to milk production, reproduction, and health. A substantial proportion of BICIs is shared with both domesticated and wild species, underlining their origin and evolutionary significance in present day cattle. In summary, our study provides valuable new resources, encompassing linked reads, de novo assemblies, NUIs, BICIs and pangenomic analyses, for future bovine pangenome research.

## Methods

### Genome sequencing of cattle breeds

Blood samples were collected from representative individuals of the Gir, Sahiwal, Tharparkar, Red Sindhi, and Kankrej breeds, in compliance with the guidelines of the Committee for Control and Supervision of Experiments on Animals (CCSEA), India. High molecular weight (HMW) DNA was extracted and 10X Genomics Chromium technology [33] libraries were prepared by AgriGenome Pvt Ltd (Genome Sequencing Company, Bangalore, India). Sequencing was subsequently performed on an Illumina HiSeq X System.

### Genome assembly

Raw data for each breed were meticulously processed for genome assembly employing the 10X Genomics Supernova assembler v2.1 (RRID:SCR_016756) [33]. The assembly process was conducted on a high-performance Dell PowerEdge R740 server equipped with 754 GB of RAM and 96 threads (48 CPUs) within a CentOS 7 Linux environment. Notably, the ’supernova mkoutput’ command was executed twice, first with the ’--style=pseudohap1’ option, producing genome assemblies representing a single haplotype for each breed. Subsequently, the command was run with the ’--style=pseudohap2’ option, which generated two assembly files representing both haplotypes within the genome of each breed [36]. These de novo assemblies serve as the foundation for downstream analyses to identify NUI as part of pangenome specific to these indigenous dairy cattle breeds.

### NUI discovery pipeline for linked-reads

The NUI discovery pipeline, as developed by Wong et al. [29], was employed in this study for the identification of NUIs in cattle. This pipeline is specifically designed for the analysis of linked-read data and has been validated using human linked-read datasets. For its execution, the NUI pipeline necessitates several prerequisites, including a reference genome, information on segmental duplications within the reference genome, and pseudohaplotypes derived from linked-read sequencing data. Additionally, the pipeline requires an alignment file in BAM format containing linked-reads aligned to the reference genome. BAM files for each linked-read dataset were generated against the Brahman reference genome using Long Ranger v2.2 (RRID:SCR_018925) [36]. Segmental duplication information for the Brahman reference genome was generated using BISER v1.4 [58].

The NUI pipeline is scripted in Bash shell and incorporates a suite of essential bioinformatics tools, including Samtools v1.2 (RRID:SCR_002105) [59], SAMBAMBA (RRID:SCR_024328) [60], BWA v0.7.15 (RRID:SCR_010910) [61], LASTZ v1.04 (RRID:SCR_018556) [62], RepeatMasker v4.1.0 (RRID:SCR_012954) [63], BEDTools v2.17.0 (RRID:SCR_006646) [64], Dustmasker [65], SAMBLASTER v0.1.24 (RRID:SCR_000468) [66], R v3.6.3 (RRID:SCR_001905) [67] and BLAST v2.9.0+ (RRID:SCR_004870) [65]. In essence, the pipeline commences by extracting unaligned reads from the BAM file using Samtools, SAMBLASTER, and SAMBAMBA, followed by trimming and quality filtering using the FASTX-Toolkit v0.0.14 (RRID:SCR_005534) [68] . These reads are then mapped individually to both pseudohaplotypes using the BWA tool. True alignments are filtered using SAMBAMBA, and genomewide coverage is calculated from the aligned files. Read clusters with coverage levels between 8X and 100X are identified using BEDTools. The pipeline proceeds to identify read clusters and extends the pseudo-haplotype sequences, expanding them by 7000 bp on each end, or until the ends of the assembled sequences, using BEDTools. These extended contigs are then aligned to the reference genome using LASTZ. The pipeline computes precise breakpoints and identifies insertions within each contig by pinpointing where the sequence alignment diverged and subsequently realigning it. Only insertional sequences with gap sizes ≥ 50 bp are retained. The outputs from the two pseudo-haplotypes are combined, and unaligned breakpoint-to-breakpoint sequences from all the contigs are extracted for subsequent analysis. These sequences are then subjected to RepeatMasker and Dustmasker analysis, with sequences containing ≥ 50 unique bases retained as NUIs. These NUIs were realigned with the reference genome using BLAST and further filtered to exclude all alignments with ≥ 95% identity and 100% coverage, while also excluding those with breakpoints overlapping assembly gaps and segmental duplications within the reference genome. Finally, NUIs identified across the five individuals are merged to create a unified and non-redundant call set, a process facilitated by the “combine_metaNUI.R” script within the pipeline, executed using Rscript. A visualization of the pipeline workflow is provided in **Fig. 1**.

### Construction of multi-assembly graph and insertion discovery

The minigraph tool was employed to construct a multi-assembly graph [21], which facilitated the discovery of insertions. The input for minigraph included the genome assembly of each breed and the Brahman reference genome. The Brahman reference genome served as the backbone, and the base alignment option was enabled (’-cxggs’) to ensure alignment consistency. Mash v2.3 (RRID:SCR_019135) [69] was used to determine the genetic distance between the Brahman genome and other cattle genomes. Based on the MASH distance calculations, the assemblies were incrementally provided to minigraph, commencing with the closest distance to the Brahman genome [23]. The output generated by minigraph was obtained in the ’.gfa’ format, which was subsequently converted to a fasta file using the gfatools v0.5 [70]. Contigs exceeding a length of 50 bp were retained as non-reference sequences, facilitating the identification of insertions within the multi-assembly graph. A visualization is provided in the flowchart in **Fig. 1**.

### Contamination screening of insertion sequences

In this study, we used an in-house, stand-alone tool known as Fasta2lineage [71], designed to identify the lineage of contigs. Fasta2Lineage utilizes the NT database of NCBI in conjunction with BLASTN for its functionality. The tool takes contig sequences in fasta format as input and conducts a similarity search against all the sequences in the database. It then identifies the best alignments and annotates their complete lineage. Contigs that do not align with any sequence in the NT database are considered novel. Fasta2lineage proves invaluable in identifying and segregating potentially contaminated sequences. For cattle assemblies, any contigs that are associated with microorganisms such as archaea, bacteria, viruses, or plants, as well as non-chordate sequences, are classified as potential contaminations.

In our study, NUIs previously identified using the NUI discovery pipeline and all insertions identified via minigraph were subjected to screening using the Fasta2Lineage tool. Sequences that were deemed novel or associated with Chordata were selected as decontaminated sequences for both datasets.

### Selection of common set of NUI and comparison with published cattle pangenomes

Cleaned contigs from NUI discovery pipeline and minigraph analysis were cross-referenced using the best bidirectional hits method with BLAST, applying stringent criteria of 95% identity and 95% query coverage. Among the selected contigs, NUIs that overlapped with insertion sequences in the minigraph dataset were selected as common and final set of NUIs.

The final set of NUIs were compared with two published pangenomes, developed by Zhou et al. [12] and Dai et al. [19]. The pangenome by Zhou et al. encompasses 22,324 contigs, collectively spanning a substantial 94 Mb, and was derived from genomic data of 898 cattle across 57 breeds. In contrast, the recently published pangenome of *Bos indicus* cattle by Dai et al. [19] was constructed from genome assemblies of 10 Chinese breeds. This recently established pangenome comprises 74,907 sequences spanning 124.4 Mb. Both sets of pangenome sequences were downloaded and subjected to comparison with the NUIs using the BLAST [65]. NUIs were considered matching when their sequences aligned with a minimum of 95% coverage and 95% identity to the published non-reference sequence.

### Identification of common insertions

To screen the presence of NUIs within the population, we performed genotyping using short-read sequence data with >30X coverage. This data was obtained by sequencing 98 individuals from five cattle breeds: Gir (20), Kankrej (19), Tharparkar (20), Sahiwal (20), and Red Sindhi (19), utilizing the Illumina HiSeqX platform. Each sample underwent preprocessing using fastp v0.23.3 (RRID:SCR_016962) [72]. Further, clean reads of each sample were mapped onto both pseudo haplotypes of each genome using BWA-MEM2 v2.2.1 (RRID:SCR_022192) [73]. The NUI discovery pipeline provided NUI fasta sequences and their coordinates on the representative pseudo haplotypes. With this information, aligned BAM files for each sample were processed to determine the depth and consensus bases at each NUI coordinate on the pseudo haplotypes using Samtools. Additionally, in-house Perl scripts [74] were employed to discern the presence or absence of NUIs within the sample. NUIs covered with at least 80% coverage and 90% identity were considered ’present,’ while those not meeting these criteria were categorized as ’absent.’ Subsequently, a genotyping matrix was generated by consolidating data from all the samples. Furthermore, NUIs with Minor Allele Frequency (MAF) <5% across all samples were excluded, and the remaining NUIs were considered as the final set of *Bos indicus* Common Insertions (BICIs) variably present in desi breeds.

### Principal Component Analysis

The NUI discovery pipeline provided information for each BICI and its occurrence in the pseudohaplotypes of each genome. From the NUI discovery pipeline output, we selectively extracted the final set of NUIs and their corresponding occurrence information. This information was used to create a BICIs occurrence matrix, which served as input for the Principal Component Analysis (PCA). The BICIs occurrence matrix was generated using a Perl script [74], with matrix elements designated as 0, 1, and 2. In this matrix, 0 represents the absence of BICI, whereas a value of 1 was assigned if the BICI was present in either of the pseudohaplotypes, and a value of 2 was allocated if the BICI was present in both pseudohaplotypes. The PCA was calculated using R, and a scatterplot of PC1 vs. PC2 was generated using SRplot [75].

### Repeat and transposable element analysis of BICIs

The fasta sequences of BICIs were extracted and analyzed to determine the composition of repeats and transposable elements (TEs) using RepeatMasker v4.1.0 (RRID:SCR_012954) [63]. RepeatMasker was executed with --species cow, -xsmall, and -nolow. Within each BICI, the TE element constituting the highest percentage of that sequence was designated as the major TE.

Furthermore, we extended the BICI sequences by 300 base pairs on each side to include flanking sequences. We then employed RepeatMasker with the same parameters to assess the composition of TEs in flanking sequences. This analysis aimed to identify TEs spanning the breakpoints on either side of the BICI and to identify major TEs present in the flanking sequences.

### Identifying transcriptionally potent BICIs

We employed the gene prediction tool Augustus v3.4.0 (RRID:SCR_008417) [76] to predict protein-coding genes within the BICIs, using the options ’--singlestrand=true --genemodel=complete’ The resulting predicted protein sequences were then subjected to functional and pathway annotation.

Additionally, we conducted a thorough examination of the breakpoint locations of BICIs within the gene sequences of the Brahman reference genome. Annotated genes that coincided with BICI breakpoints were identified and extracted for analysis. For BICIs with breakpoints outside of genes, we conducted a search for the nearest gene within a 20-kilobase (20 Kb) range.

To further investigate the transcriptional potential of BICIs, we mapped RNA-Seq data from 47 samples representing various indigenous breeds, including Gir, Tharparkar, Sahiwal, and Haryana, onto the Brahman reference genome. The genome was indexed using the STAR aligner v2.7.10 (RRID:SCR_004463) [77] with the command: ’STAR --runThreadN 32 --runMode genomeGenerate --genomeDir ./ --genomeFastaFiles

./GCF_003369695.1_UOA_Brahman_1_genomic.fna --sjdbGTFfile GCF_003369695.1_UOA_Brahman_1_genomic.gtf --sjdbOverhang 100.’

We aligned all RNA-Seq reads to this indexed genome, one sample at a time, using the command: ’STAR --runThreadN 40 --outReadsUnmapped Fastx --genomeDir ./assembly/ --outFileNamePrefix ./dis_04R --outFilterMultimapNmax 100000 --outSAMunmapped Within KeepPairs --limitOutSAMoneReadBytes 1000000 --readFilesCommand zcat --readFilesIn read1.fq.gz read2.fq.gz.’ This process generated separate R1 and R2 files in FASTQ format for the unmapped reads. Subsequently, we collected the unmapped reads from all 47 samples and merged these FASTQ files into single read1 and read2 files. We then realigned these merged unmapped reads to the Brahman reference genome without supplying the GTF file. Again, we collected all the unmapped reads in Fastq format and aligned them to BICI with flanking sequences extended by 300 bp on both ends. We generated an index of flanked BICIs and aligned the unmapped reads to these indexed BICIs using parameters as previously mentioned, with the exception of not providing a GTF file. We filtered the resulting alignment file in SAM format based on the following criteria: (a) both reads in a read pair were mapped. (b) read pairs were mapped in the correct orientation and with the correct insert size. (c) reads had alignment scores ≥ 140, equivalent to four total mismatches for a read pair. (d) reads had a mapping quality score of 255, indicative of unique mapping according to the STAR scoring scheme.

The filtered alignment file was converted into a coordinate-sorted BAM file and processed using Stringtie v2.1.1 (RRID:SCR_016323) [46] to identify novel transcripts within the BICI sequences. The command ’stringtie -o nui_mapped_STAR.gtf filtered_sorted_STAR.bam -p 10’ was used. Transcripts that fully resided within the 300 bp flanking sequences were discarded. This analysis enabled us to identify all novel transcripts and their corresponding transcribed BICIs.

### Functional analysis of genes with BICIs

GO enrichment analysis was performed on non-redundant set of genes having BICIs using the clusterProfiler package (RRID:SCR_016884) [78] from the Bioconductor in R (RRID:SCR_006442) [79]. A significance threshold of p-value ≤ 0.05 (FDR by Benjamini–Hochberg) was applied to identify significant enriched terms. The resulting GO IDs from the clusterProfiler package, along with their respective p-values, were then subjected to REVIGO (RRID:SCR_005825) [80] to eliminate redundant GO categories from all enriched terms. Finally, the results of all GO categories, in conjunction with the clusterProfiler package, were visualized using a Python-based tool CirGO [81].

To establish a connection between BICIs and major QTL, non-redundant genes featuring BICIs were aligned with the publicly accessible cattle QTL Database [82]. The mapping process involved extracting non-redundant genes with BICIs and conducting a BLAST search against the CDS annotated in ARS_UCD 1.2 assembly to identify orthologous genes. Subsequently, these orthologous genes were cross-referenced with the QTL database, allowing for the characterization of their associations with specific traits.

### Evolutionary analysis of the BICIs

To investigate the presence of BICIs in related species, we first aligned BICIs to the genomes of sister species within the Bos genus, which includes *Bos taurus* (taurine cattle) [22], *Bos gaurus* (gaur) [83], *Bos grunniens* (domestic yak) [84], *Bos frontalis* (gayal) [85], and *Bos mutus* (wild yak) [84]. In a subsequent analysis, we extended our alignment to species of bovidae family which includes *Bubalus bubalis* (buffalo) [86], *Tragelaphus oryx* (eland) [87], *Bison bison* (American bison) [88], *Capra hircus* (goat) [89], and *Ovis aries* (sheep) [90]. Reference genome sequences for each of these species were obtained from NCBI, and BICIs were aligned to their respective reference genomes using BLAST. Alignments with a minimum of 95% identity and 95% query coverage were considered genuine matches. Distribution of BICIs across species were plotted using jvenn (RRID:SCR_016343) [91].

## Additional Files

Additional File1: BICI fasta file

Additional File2: Final NUI set fasta file

Additional File3: Minigraph pipeline fasta file

Additional File4: NUI discovery pipeline fasta file

## Data availability

The Linked Read data generated and reported in this article are accessible from the Indian Biological Data Centre (IBDC) with the INSDC accession number mentioned in the (**Supplementary Table S9**) Additionally, the Supernova assemblies generated from the raw Linked Reads is also available on IBDC. The Illumina sequencing data used in this study have been submitted to IBDC, and all accession numbers are detailed in (**Supplementary Table S10**). RNAseq data with all accession numbers detailed in (**Supplementary Table S11**). Given that IBDC is part of the INSDC, all the data can be accessed from the NCBI as well.

## Competing Interest

The authors declare that they have no competing interests.

## Funding

The Linked-Read data were generated in the project “Genomics for Conservation of Indigenous Cattle Breeds and for Enhancing Milk Yield, Phase-I” [BT/PR26466/AAQ/1/704/2017], funded by the Department of Biotechnology (DBT), India. Transcriptomics data were generated in the project “Identification of key molecular factors involved in resistance/susceptibility to paratuberculosis infection in indigenous breeds of cows” [BT/PR32758/AAQ/1/760/2019], which was also funded by Department of Biotechnology (DBT), India.

## Authors’ contributions

S.A., B.D.R., S.N.R. and S.S.M designed the study. S.S.M, R.K.G and S.A. facilitated Sample collections and sequencing. A.S. and N.K.P. assembled the genomes and established the contamination screening pipeline. S.A. and A.S. analyzed the data for NUIs identification through NUI discovery pipeline, performed minigraph analysis, Genotyping of NUIs in the population, PCA, TE analysis, GO analysis and evolutionary analysis. M.N. performed QTL analysis. R.K.G., C.P.V.T., B.D.R and S.A. drafted the manuscript. S.S.M., S.N.R. and C.P.V.T. edited the manuscript. All authors reviewed the final manuscript before submission.

## Acknowledgements

The authors sincerely express their gratitude to the Department of Biotechnology (DBT), Ministry of Science, New Delhi, India for providing financial support. Additionally, the authors extend their appreciation to the National Institute of Animal Biotechnology (NIAB) for offering invaluable support throughout the execution of the study. S.A. specifically acknowledges and appreciates the support received from Dr. G. Taru Sharma, Director, NIAB. M.N., C.P.V.T., and B.D.R. were supported by the appropriated project 8042-31000-112-000-D, “Accelerating Genetic Improvement of Ruminants Through Enhanced Genome Assembly, Annotation, and Selection” of the USDA Agricultural Research Service. Any mention of trade names or commercial products is solely for the purpose of providing specific information and does not imply recommendation or endorsement by the U.S. Department of Agriculture. The USDA is an equal opportunity provider and employer.

## Abbreviations

BAM: Binary Alignment Map
BICI: *Bos indicus* Common Insertions
bp: base pairs
CDS: Coding sequence
FDR: False Discovery Rate
GB: Gigabyte
Gb: Gigabase
gfa: graphical fragment assembly
GO: Gene Ontology
Kb: Kilobase
MAF: Minor Allele Frequency
Mb: Megabase
NCBI: National center of Biotechnology
NUI: Non-reference Unique Insertion
PCA: Principal Component Analysis
SNP: Single Nucleotide polymorphism
SV: Structural variation
TE: Transposable Element
QTL: Quantitative trait loci

## Supplementary Figure Legend

**Figure S1:**
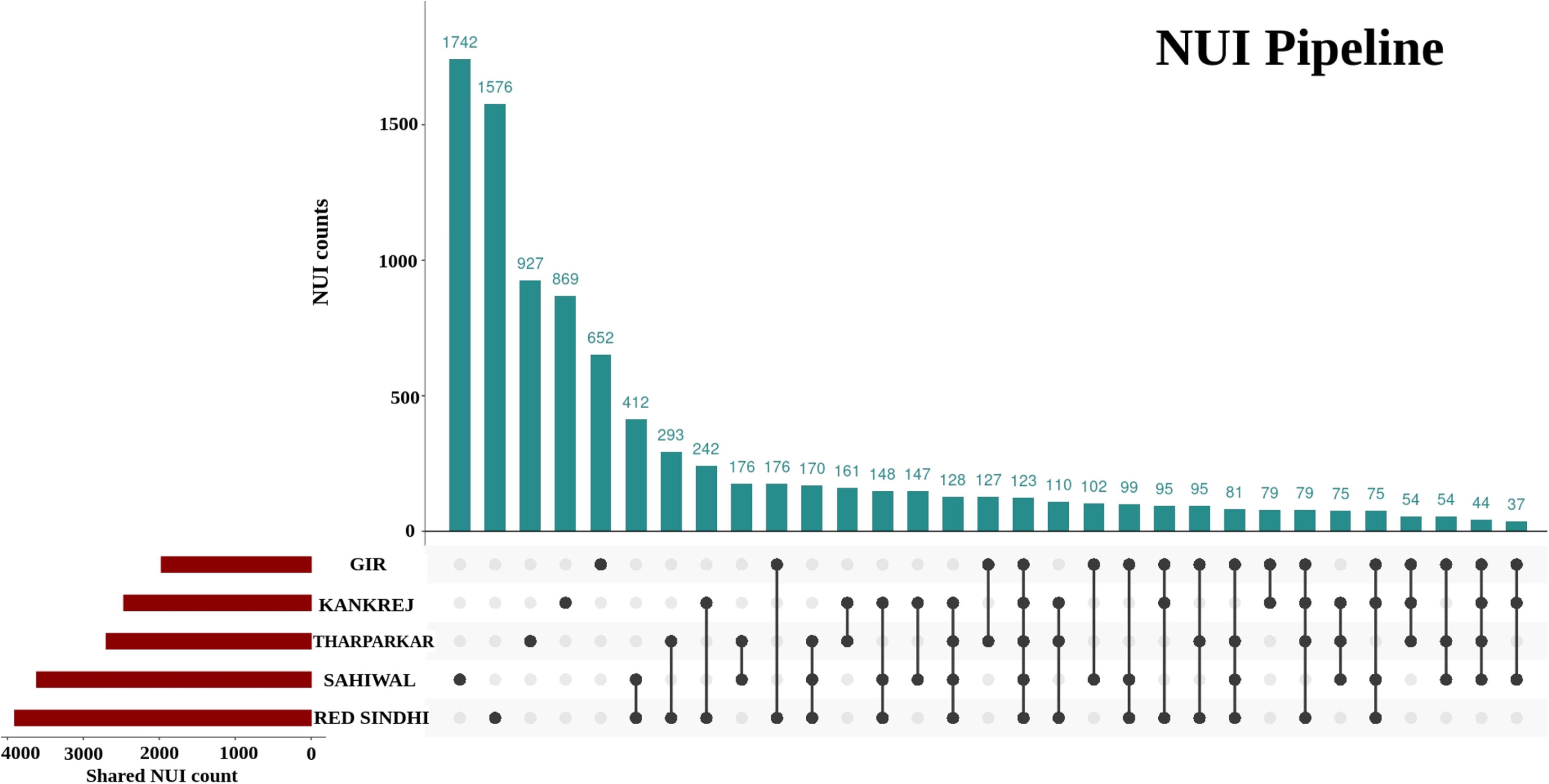
Upset plot for the shared Non-Unique Insertions (NUIs) among five *Bos indicus* breeds from NUI discovery pipeline. Each vertical bar in the chart signifies the number of NUIs shared by the corresponding combination of breeds listed below. The horizontal bar chart represents the count of these shared NUIs.

**Figure S2:**
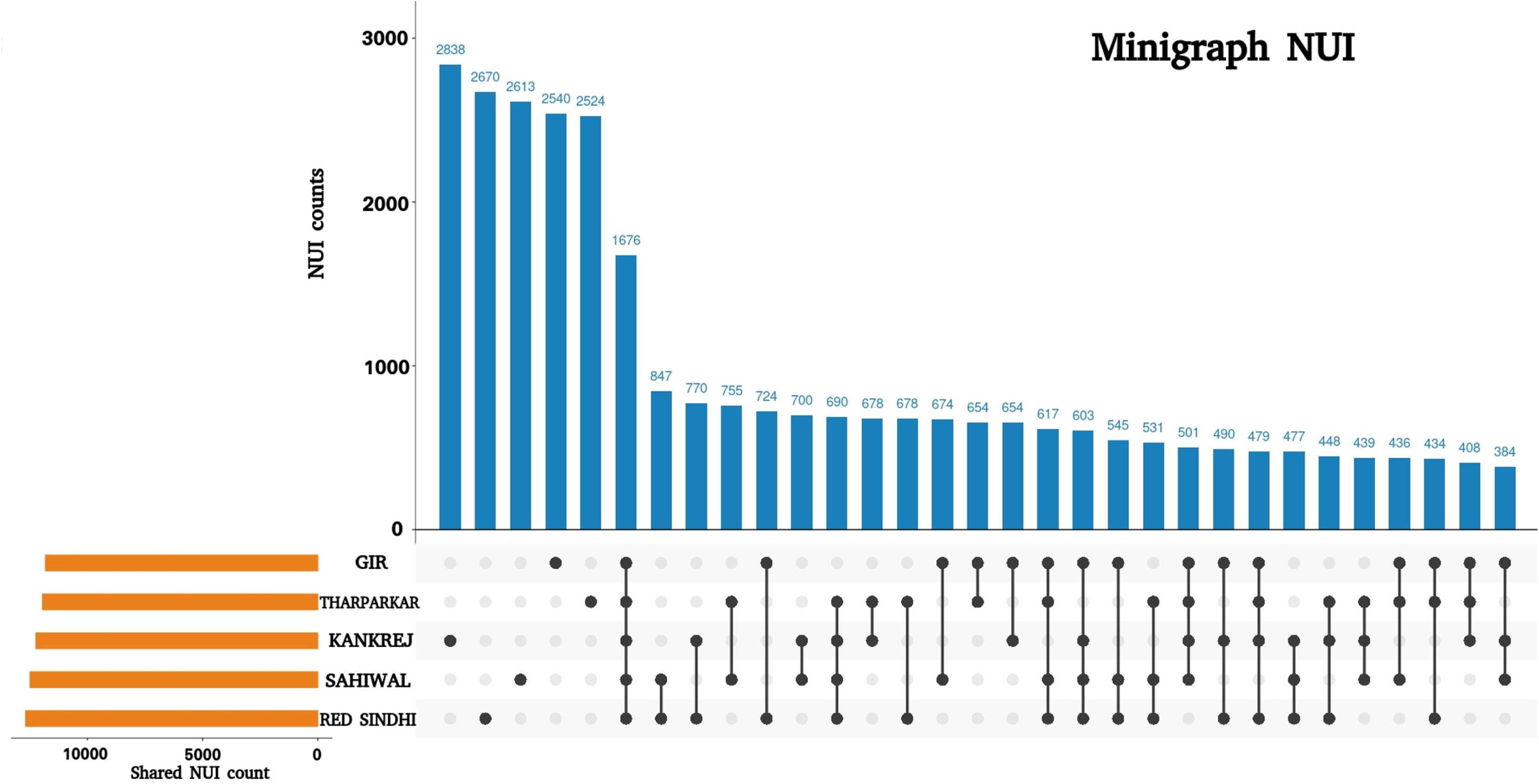
Upset plot for the shared Non-Unique Insertions (NUIs) among five *Bos indicus* breeds from minigraph pipeline. Each vertical bar within the chart denotes the quantity of NUIs shared by the respective combination of listed breeds below. The horizontal bar graph indicates the tally of these shared NUIs.

**Figure S3:**
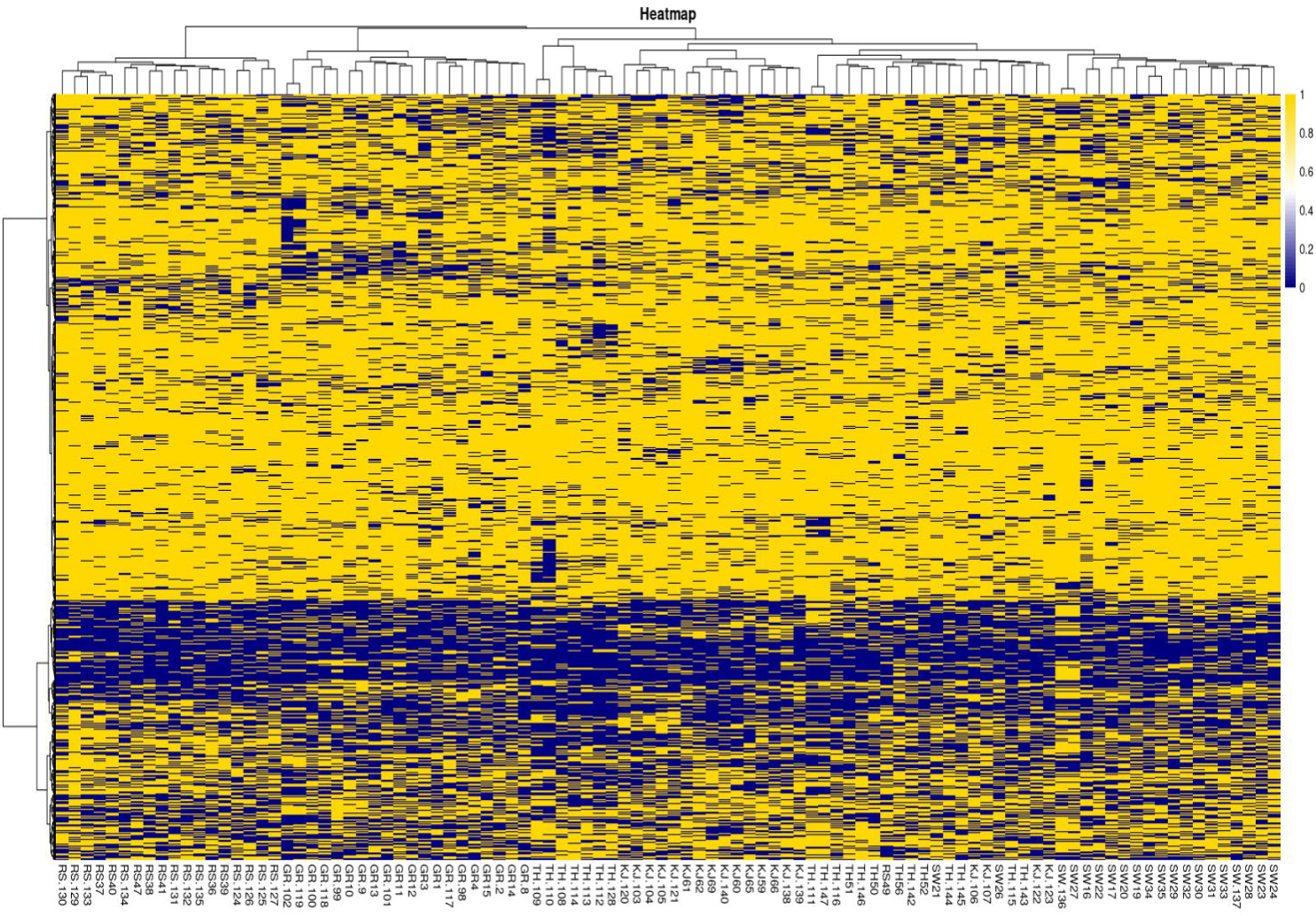
Heatmap of presence and absence of Non-Unique Insertions (NUIs) across the 98 samples. The presence of NUIs is depicted in yellow, while their absence is represented by navy blue color. Furthermore, the breeds are clustered based on Euclidean distance

**Figure S4:**
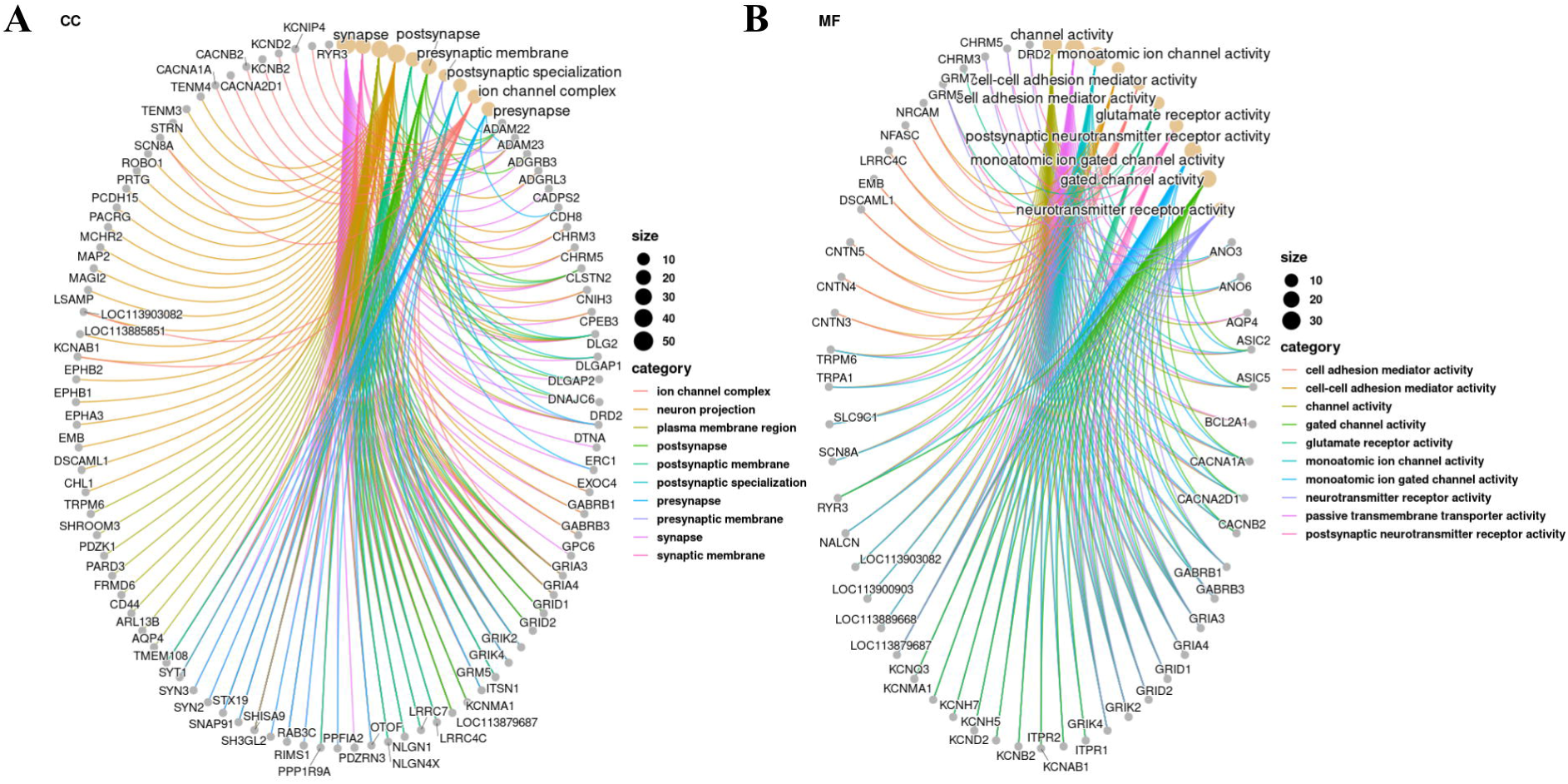
GO annotation of BICIs. **(A)**The Cnet plot displays the enriched terms and genes associated with cellular components. **(B)** The Cnet plot illustrates the enriched terms and genes pertaining to molecular function.

## Notes

### Competing Interest Statement

The authors have declared no competing interest.

